# Genome-scale metabolic models consistently predict *in vitro* characteristics of *Corynebacterium striatum*

**DOI:** 10.1101/2023.04.28.538764

**Authors:** Famke Bäuerle, Gwendolyn O. Gusak, Laura Camus, Simon Heilbronner, Andreas Dräger

## Abstract

Genome-scale metabolic models (GEMs) are organism specific knowledge bases which can be used to unravel pathogenicity or improve production of specific metabolites in biotechnology applications. The present work combines *in silico* and *in vitro* approaches to create and curate strain-specific genome-scale metabolic models of *Corynebacterium striatum*. Approaches towards modeling rarely studied organisms are scarce. We introduce a cost-effective and easy experimental protocol which can be adapted to other organisms as well. Furthermore, the comparability of growth kinetics and *in silico* growth rates is discussed. This work introduces five newly created strain-specific genome-scale metabolic models (GEMs) of high quality, satisfying all contemporary standards and requirements. All these models have been benchmarked using the community standard test suite Metabolic Model Testing (MEMOTE) and experimentally validated by laboratory experiments conducted specifically for this purpose. For the curation of those models, the software infrastructure *refineGEMs* was developed to work on these models in parallel and to comply with the quality standards for GEMs. The model predictions were confirmed by experimental data and a new comparison metric based on the doubling time was developed to quantify bacterial growth. Future modeling projects can rely on the proposed software, which is independent of specific environmental conditions. The validation approach based on the growth rate calculation is now accessible and closely aligned with biological questions. The curated models are freely available via BioModels and a GitHub repository and can be used. The open-source software refineGEMs is available from https://github.com/draeger-lab/refinegems.

## 1 Introduction

Human life and substantial cultural achievements depend on prokaryotes. While a minority of bacterial species cause invasive diseases, humans could not exist without beneficial microbes colonizing the gut and the body surfaces. Additionally, the metabolic capacities of microbes flavor and preserve our food, detoxify the environment, or allow the production of antibiotics to treat infections. Without the ability to identify and grow microbes under defined conditions within laboratory surroundings, many modern comforts would be unthinkable. Remarkably, we are only beginning to appreciate the enormous diversity of bacterial life, and even with state-of-the-art techniques to grow bacteria, the vast majority remains uncultivated^1^.

As a huge metabolic capacity is slumbering within these “unculturable” bacteria, efforts for their cultivation should be intensified^2, 3^. Computer modeling of the nutritional needs of microorganisms holds great promise in this regard. The importance of computational modeling in biology has steadily increased over the last decades^4^. In particular, genome-scale metabolic models, so-called GEMs, have proven beneficial in numerous application areas due to their facile mathematical manageability and predictive power^5^.

Among other benefits, GEMs permit predicting an appropriate media composition that should allow proliferation for any organism. However, up to date, this potential seems hardly exploited as it needs combined efforts of bioinformaticians to make predictions and of microbiologists to test the same systematically. Most published models lack sufficient experimental validation, or the designators of the components contained in the models only allow preliminary conclusions to be drawn about tangible cell components. These shortcomings reduce the validity or interpretability of the predicted results. Only if modelers and experimenters work closely together can they check predictions made by the model and, if necessary, change the structure of the model or the experimental setup to gradually arrive at a meaningful representation of reality in the model that yields reliable results^6^.

Herein we tested the ability of *de novo* created GEMs to predict *in vitro* growth characteristics of *Corynebacterium striatum*. At the time of writing, no genome-scale systems biology models are available that characterize this species. *C. striatum* is a Gram-positive, non-sporulating rod discovered in the early 20th century. It was considered a commensal within the healthy human skin microbiota for a long time^7^, and its pathogenic properties remained unknown. With increasing awareness of nosocomial infections, *C. striatum* was identified as a cause of diseases, particularly for immunocompromised patients. It may cause several diseases, including Chronic Obstructive Pulmonary Disease, COPD, or pneumonia^8^. In addition to the human respiratory tract, longstanding open wounds belong to its habitat^9^, leading to prolonged hospitalizations^10^. Compared to other Gram-positive members of the skin flora, *C. striatum* is particularly resistant to several antibiotics^7^, including ampicillin, penicillin, and tetracycline^9^. For these reasons, *C. striatum* constitutes an ideal example case for benchmarking and improving a systematic and semi-automatic modeling environment.

The individual work steps required to reconstruct highquality models have been excellently characterized in the literature in numerous examples^11–14^. However, a substantial obstacle in this endeavor is the enormous complexity and the sheer number of those steps, which Thiele and Palsson put at 96 repetitive steps^15^. In addition, this requires knowledge from many orthogonal scientific disciplines ranging from (bio)chemistry, biophysics, and bioinformatics to mathematics and their application in bioengineering and, depending on the use case, various fields such as microbiology, oncology, or biotechnology. Working with the underlying file formats and data standards also requires specialized knowledge in particular areas. It is easy to see why the modeling work is neither intuitive nor generally accessible. Fundamental programming skills are required to examine the resulting models just rudimentarily.

The file format Systems Biology Markup Language (SBML) Level 3^16^ is most widely used to encode systems biology models^17^. However, a particular extension, the Flux Balance Constraints (FBC) package^18^, is required to store constraint-based genome-scale models in it. Other packages open up additional features of the SBML^19^ format to users, such as linking metabolic maps directly to the computational model^20, 21^, offering a wide range of visualization capabilities^22, 23^.

Such metabolic maps are often drawn using web-based programs such as Escher^24^ or Newt^25^, which are compatible with the Systems Biology Graphical Notation (SBGN) standard^26, 27^ and generate unambiguously represented biological network maps. The computational model can be created and edited using various programs, depending on the type of modeling. Constraint-Based Reconstruction and Analysis (COBRA) programs such as COBRApy^28^ (for Python users) or COBRA Toolbox^29^ (for Matlab users) have proven particularly useful for genome-scale models. An initial model is, in turn, first created with programs such as CarveMe^30^. For the publication of models, the BioModels database is usually available^31^.

This short list is not exhaustive but should illustrate that modeling work is often discontinuous because numerous different programs are required in combination. In addition, the modeler needs to think about a suitable structure for versioning their models and save it in a way that is understandable to others. Many steps also require manual revision, sometimes done directly in the SBML^19^ files or using the command line and highly specific scripts. Much of the work required for modeling may be similar in its outcome but repetitively and independently developed. Not only does the resulting redundancy lead to endless hours of avoidable programming work. It also results in projects that are incompatible with each other but similar in their effect^32^.

The fewer users test a particular software project, the higher the probability of finding previously undiscovered programming errors. Ultimately, these also affect the model quality. Since each developer typically places the aspects relevant to its respective project in the foreground and cannot cover the curation of the models in sufficient depth, it can be easily explained why frequently certain aspects fall by the wayside: either due to a lack of time or the sheer overwhelming nature of the overall complexity.

The present work addresses such problems by providing a unified directory structure and a collection of directly executable programs within a Git-based version control system. To this end, the authors use their experience gained in numerous previous projects on systems biology modeling^12, 14^, software development^33, 34^ and laboratory work^35, 36^ to precisely reconstruct and experimentally validate multiple strains in parallel using currently common standard operating protocols for a bacterium that remains to be studied in systems biology. In this work, we constructed GEMs of five *C. striatum* model strains and used them to predict growth characteristics under defined nutritional conditions. These predictions were then tested in the laboratory. Interestingly, we found that the *in vitro* and *in silico* data for *C. striatum* largely overlap. In addition to the comparison between laboratory and *in silico* strain behavior, a program to enhance the creation of high-quality models is presented.

## 2 Materials and Methods

### 2.1 Genome Sequences

Both the National Center for Biotechnology Information (NCBI) and PATRIC were used to search for complete genome sequences of *C. striatum* strains. On NCBI the genome assembly ASM215680v1 of *C. striatum* KC-Na-01 with the accession reference `GCF_002156805.1` was the most prominent. This sequence was used to create a first GEM. This strain was not available to order from the *Deutsche Sammlung von Mikroorganismen und Zellkulturen* (DSMZ, translates to German Collection of Microorganisms and Cell Cultures). Thus, four additional strains with BacDrive sequence information were obtained from DSMZ (Table 1). All sequences were downloaded from NCBI (Table 1).

**Table 1.** *C. striatum* strains used in this study, with the corresponding DSM-number, NCBI accession number, and identifier. “Short ID” is the name used in this manuscript. The “Model ID” contains the final ids of the models uploaded to BioModels where it is accessible using the identifier (ID) in the rightmost column. TS denotes the type strain of *C. striatum*.

| Name | DSM | NCBI | Short ID | Model ID | BioModels ID |
| --- | --- | --- | --- | --- | --- |
| FDA-ARGOS_1054 | 20668 | CP066290 | TS | iCstr1054FB23 | MODEL2304270001 |
| FDA-ARGOS_1197 | 45711 | CP069514 | 1197 | iCstr1197FB23 | MODEL2304270003 |
| FDA-ARGOS_1115 | 7184 | CP068158 | 1115 | iCstr1115FB23 | MODEL2304270002 |
| FDA-ARGOS_1116 | 7185 | CP068157 | 1116 | iCstr1116FB23 | MODEL2304270004 |
| KC-Na-01 | - | - | KC | iCstrKCNa01FB23 | MODEL2304270005 |

### 2.2 Existing Model Assessment

One GEM for *C. striatum* exists in the Virtual Metabolic Human (VMH) database. The VMH offers GEMs of organisms interesting for human microbiomes. Most models within the VMH database are used as a basis to build simulatable microbiomes. Additionally, all VMH GEMs use specific entity IDs of which some overlap with biochemically, genomically, and genetically structured (BiGG)^37^ IDss. The one existing GEM was assessed with MEMOTE and manually evaluated by looking at the Extensible Markup Language (XML) file.

### 2.3 Draft Models

Draft models were created from the genome sequences annotated by the NCBI Prokaryotic Genome Annotation Pipeline using CarveMe^30^. We used version 1.5.1 of the package, which we installed via `pip` on a MacBookPro running macOS Monterey version 12.3.1. CarveMe was run on the protein FASTA files of the sequences with the `fbc2` flag. Fbc2 refers to the SBML Level 3^16^ package “Flux Balance Constraints”^18^ which extends models written in SBML^19^ by structures that enable flux bounds and optimization functions. These structures are necessary for growth simulations with flux balance analysis (FBA) which were used to compare the models to laboratory results.

### 2.4 Automated Polishing

The drafts were polished with ModelPolisher^38^ and with the script 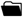 `polish.py` available with the refineGEMs toolbox (see Section 3.3). Automated polishing included moving entries from the notes section to the annotation section of an entity, annotating all entities with their respective BiGG^37^ ID as identifiers.org link, and setting the models parameters to `mmol_per_gDW_per_h`. In addition, the GeneProducts were polished using refineGEMs: They were annotated with the NCBI Protein ID and renamed with the respective name indicated on NCBI.

### 2.5 Semi-automated Curation

RefineGEMs was used to add charges to metabolites that had no denoted charge, some of the missing charges were extracted from the model of *Pseudomonas putida* KT2440 with the ID *i*JN1463. RefineGEMs was used to apply the SBOannotator tool^39^ which allows for automated Systems Biology Ontology (SBO)^40^ term annotation and specialization. The Python module `MassChargeCuration`^41^ was used to further correct the charges and masses of the metabolites. RefineGEMs was used to synchronize annotations of metabolites in different compartments.

### 2.6 Manual Curation

Manual curation was first focused on cleaning residues left from the automated draft reconstruction: Duplicate reactions and metabolites with different IDs were identified using MEMOTE ^42^ and subsequently removed. Metabolites with only a few or no annotations were researched manually in multiple databases and annotated based on the results. Network gaps were identified by drawing pathways as Escher maps^24^ and comparing those to Kyoto Encyclopedia of Genes and Genomes (KEGG) pathway maps^43^.

### 2.7 Model Quality Assessment

The MEMOTE score^42^ serves as a tool for researchers to quickly evaluate the completeness and accuracy of GEMs and to compare models across different organisms and strains. It also helps to identify potential gaps in the models, making it easier to improve them in the future.

### 2.8 *In silico* growth rates

Growth rates and, thus, doubling times were determined by FBA. We used the routine implemented in refineGEMs which is based on the COBRApy optimize function. This maximized the flux through the biomass objective function (BOF). It assumes that the goal of any organism is to increase its biomass toward cell division. The returned objective value was then interpreted as growth rate in mmol*/*(g_DW_ · h). The doubling time or generation time *T*_*d*_ in minutes is then calculated by the formula

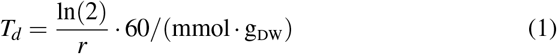

with *r* being the growth rate or objective value extracted from the FBA. All used media formulations can be found in the 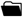 `sbo_media_db.sql` in refineGEMs.

### 2.9 Growth Rate Comparison

Growth rates from simulations with COBRApy are given in mmol*/*(g_DW_ · h), which can be used to calculate doubling times using Equation (1). To ensure robust comparison, we chose to extract doubling times from experimental growth curves by fitting a logistic equation to the data and using the rate at the inflection point as the growth rate in optical density (OD) per time from which we can calculate the doubling time. This approach was taken for all comparisons.

### 2.10 *In vitro* growth phenotypes

All media compositions are available in the Supporting Information. Cultures were grown in Tryptic Soy Broth (TSB) overnight at 37 ^*°*^C and 150 rpm. After 10 min of centrifugation at 4,000 rpm, the remaining pellets were resuspended in the medium of interest. The samples were inoculated at an OD of 0.1. The OD _600 nm_ (OD600) was measured at *t* = 0 h and at *t* = 24 h. Fold changes were calculated by dividing OD600 (*t* = 24 h) by OD600 (*t* = 0 h).

### 2.11 *In vitro* growth rates

For each of the four strains, one colony grown on Tryptic Soy Agar (TSA) Blood plates was inoculated in 10 mL of TSB. Precultures were incubated overnight at 37 ^*°*^C and 150 rpm and diluted to an OD of 0.05 in TSB, Brain-Heart-Infusion Broth (BHI), Roswell Park Memorial Institute cell culture medium 1640 (RPMI), Lysogeny Broth (LB), M9 minimal medium (M9), and *Corynebacterium glutamicum* minimal medium version 12 (CGXII), Table S1. 150 µL of the suspensions was distributed in a Greiner Bio-One 96 flat bottom well plate and incubated at 37 ^*°*^C for 24 h in the BioTek Epoch 2 Microplate Reader. OD600 was monitored every 15 min after agitation at 600 rpm. All experiments were performed using three technical replicates per plate and at least three biological replicates.

### 2.12 Growth data analysis

Growth data were extracted from the plate reader in a Microsoft Excel^®^ file and were then read using pandas^44^. The logistic equation for growth was fitted using `curve_fit` from `scipy.optimize`^45^. This uses a non-linear least squares approach to fit Equation (2) to the growth data. Then the growth rate *r* can be extracted, and the doubling time or generation time *T*_*d*_ is then calculated by *T*_*d*_ = ^ln 2^*/*_*r*_. A logistic function that results from a logistic model of bacterial growth is shown on Equation (2):

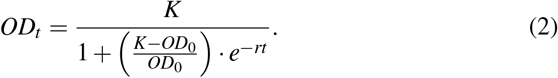

Here, *OD*_*t*_ is the *OD* at time *t, K* is the asymptote (usually the maximum *OD*), *OD*_0_ is the OD at time 0, *d* refers to the displacement along the *x*-axis. Multiple columns with *OD* values are possible. Coming from the plate reader, technical replicates were averaged before fitting. The wells representing the blank were also averaged and subtracted from the averaged technical replicates.

For the logistic fit, `curve_fit` was always initialized with `p0 = np.asarray([0.2, 0.05, 0.05])`, the function is defined in Python as

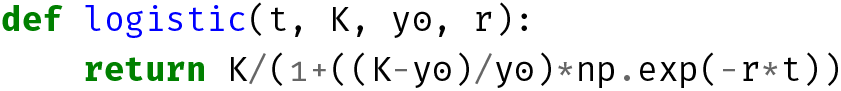

### 2.13 Model monitoring

During the tests on different media, the models were continuously monitored, and simulations were run to obtain missing metabolites that might help to recover growth on the minimal media of interest. We noticed that the models had no exchange reactions for sodium which was changed to reflect the ability of sodium uptake of the respective organisms.

## 3 Results

### 3.1 Construction of five strain-specific GEMs

#### 3.1.1 *Available model and strains of* C. striatum

To the authors’ best knowledge, no other manually curated high-quality GEM of *C. striatum* is currently available. However, within the gut microbiota resource of the VMH database^46^, a model for the type strain ATCC 6940 exists^47^.

The VMH model for *C. striatum* has a MEMOTE score of 88 %, even though it contains erroneous annotations with NaN IDs and incorrect InChIKeys. All the GeneProduct objects lack annotation, and some of the reactions were only annotated with their respective SBO terms. Thus, within this study, we created manually curated high-quality GEMs for *C. striatum*.

We decided to work with multiple strains to get a comprehensive picture of the strain-specific properties of *C. striatum*. Apart from the type strain FDAARGOS_1054 / ATCC 6940 (type strain (TS)), we used three more strains, namely, FDAARGOS_1197 (1197), FDAARGOS_1115 (1115) and FDAARGOS_1116 (1116), whose genome sequences have been completely assembled. In addition to these four strains, we also investigated the strain KC-Na-01 (KC), which is well characterized by the KEGG database, but unlike the other four strains, it is not available from the DSMZ. For this reason, for strain KC, we exclusively performed an *in silico* analysis.

The strain-specific models were named following the guidelines given by Carey et al. (2020). However, as there were no indications for naming strain-specific models we decided to elongate the species indicator by a strain indicator (in our case the FDA-ARGOS ID). As the iteration identifier the author’s initials (FB) and the year (23) were added.

#### 3.1.2 Characteristics of the models created in this study

The models created in this study are composed of 1,053 to 1,382 metabolites, 1,541 to 2,002 reactions, and 719 to 772 genes. This is within the usual range of bacterial GEM compare, e.g., with 1,183 genes and 2,276 reactions 49). Figure 1A and Table S2 show that the model of strain TS has the biggest scope with respect to the number of entities that it holds. All GEMs have a metabolic coverage above 2 %. Usually, models with a high level of modeling detail will have a metabolic coverage above 1 %, which highlights the degree of detail of our models. All MEMOTE scores are above 84 %, which is higher than all scores of models evaluated for the MEMOTE meta-study in 2020^42^ (compare SI Figure 30 of that study). All entities that are present in the models are annotated both with Uniform Resource Identifiers (URIs) and SBO terms. We used ten different databases for metabolite annotations, eight for reaction annotations, and two databases to annotate protein-encoding sequences which are added as Gene Protein Reactions to the models. KEGG metabolic pathways were added to all models. The models were checked for energy-generating cycles, which were eliminated, and orphaned metabolites were connected to the network. All models are stoichiometrically consistent.

**Figure 1.**
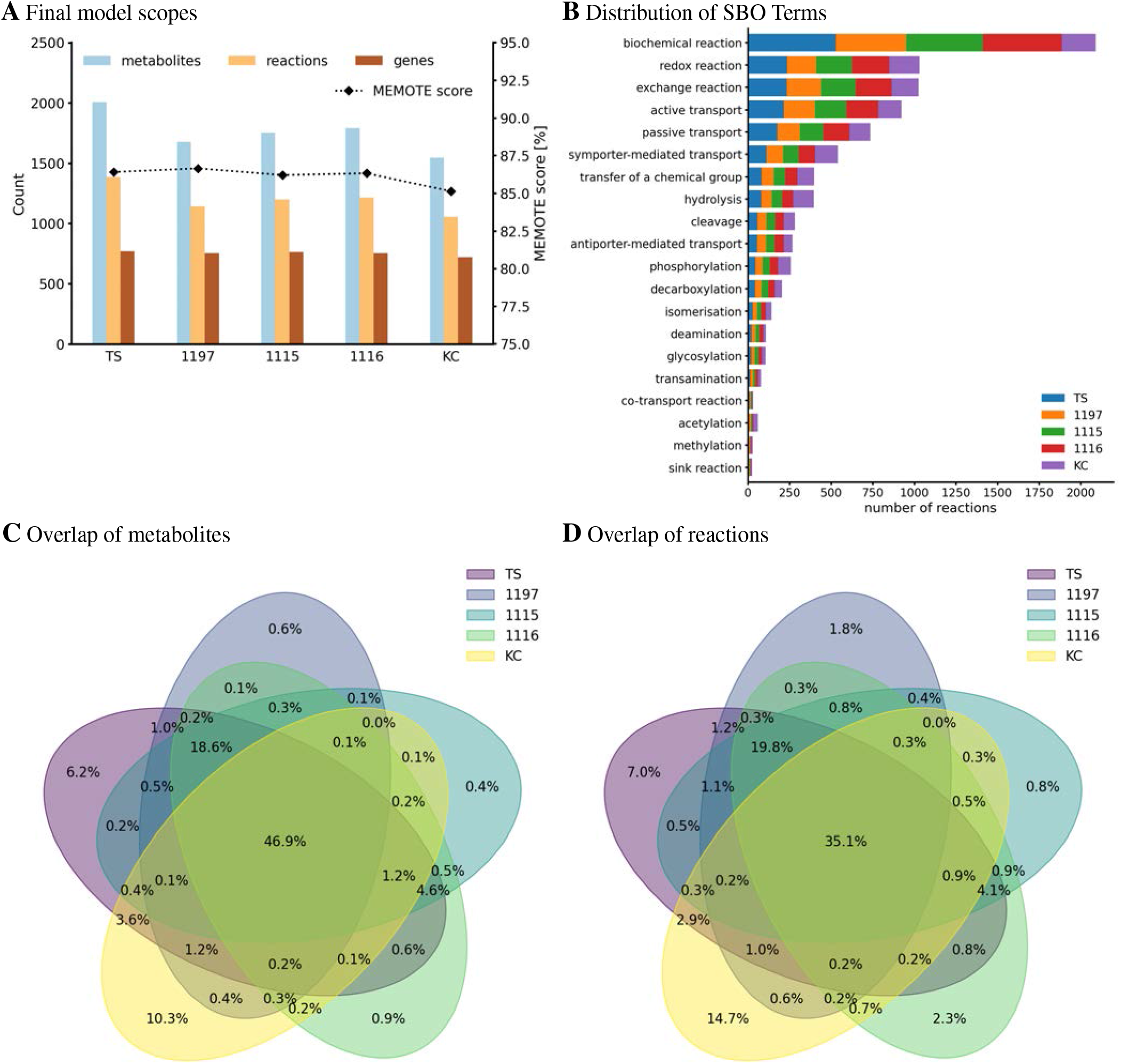
**A) Scopes of all curated *C. striatum* models**. Number of reactions, metabolites, and genes are read on the left y-axis, MEMOTE scores are read on the right y-axis. This score is based on standardized and community-maintained metabolic tests for quality control and quality assurance of GEMs. **B) Classification of reaction types by assigned SBO terms**. SBO terms were annotated using the SBOAnnotator^39^ tool and thus are specialized. The distribution of different reactions is shown with stacked bars which represent the number of reactions that are classified with that SBO term. **C) and D) Strain comparison based on the metabolic reconstructions**. Venn diagrams showing the overlap of all metabolites (C) and all reactions (D) of the curated models in this study. These diagrams were created based on the metabolite/reaction identifiers using the pyvenn Python module.

Reactions can be classified into different types using the SBO terms. SBO analysis showed that all four models follow a similar reaction type distribution (Figure 1B), suggesting vastly overlapping metabolic capacities of the strains.

The metabolic reconstructions were used for a detailed comparison of the five strains. This showed an overlap of 47 % for all metabolites and an overlap of 35 % for all reactions across all models (Figure 1C and Figure 1D). With 168 and 391 unique metabolites and reactions, respectively, the strain KC showed higher divergence compared to the other models. In comparison, strain 1115 showed 6 and 21 unique metabolites and reactions, respectively. This suggests a reduced metabolic fitness compared to strain KC.

### 3.2 Experimental validation of the strain-specific GEMs

GEMs can be used to predict growth characteristics of bacterial species under defined nutritional conditions^14^. This can be useful for laboratory experimentation, especially when little to no *in vitro* data for the microorganism of interest is available. However, the accuracy of GEMs in predicting *in vitro* characteristics is frequently unclear. Therefore, we decided to compare model predictions to *in vitro* growth characteristics under various nutritional conditions.

#### 3.2.1 Growth characteristics under varying nutritional conditions

We used the complex, nutritionally rich medium LB^50^ for which a nutritional composition to be used in metabolic modeling is available^30^. However, this medium contains the complex component yeast extract, making the precise composition of the medium *in vitro* unclear. As defined media, we used RPMI^51^, M9^52, 53^, and CGXII^54–56^ since their precise chemical composition is known.

*In vitro* experiments allow for different experimental approaches. Besides changing incubation temperature, bacterial cultures can be grown in flasks with or without a baffle whose volumes can be adapted from microliters to liters. Alternatively, cultures can be grown in microtiter formats (100 to 500 µL*/*well) or even in continuous culture systems where a constant refreshment of medium with an inflow of metabolites can be used. It is well known that varying growth conditions impact the achieved cell densities and growth rates significantly. However, the influence such conditions have on, e.g., gas exchange rates, are difficult to reflect within GEMs.

We, therefore, decided to test the accuracy of the model to predict the growth characteristics of the strains under two distinct growth conditions (Figure 2).

**Figure 2.**
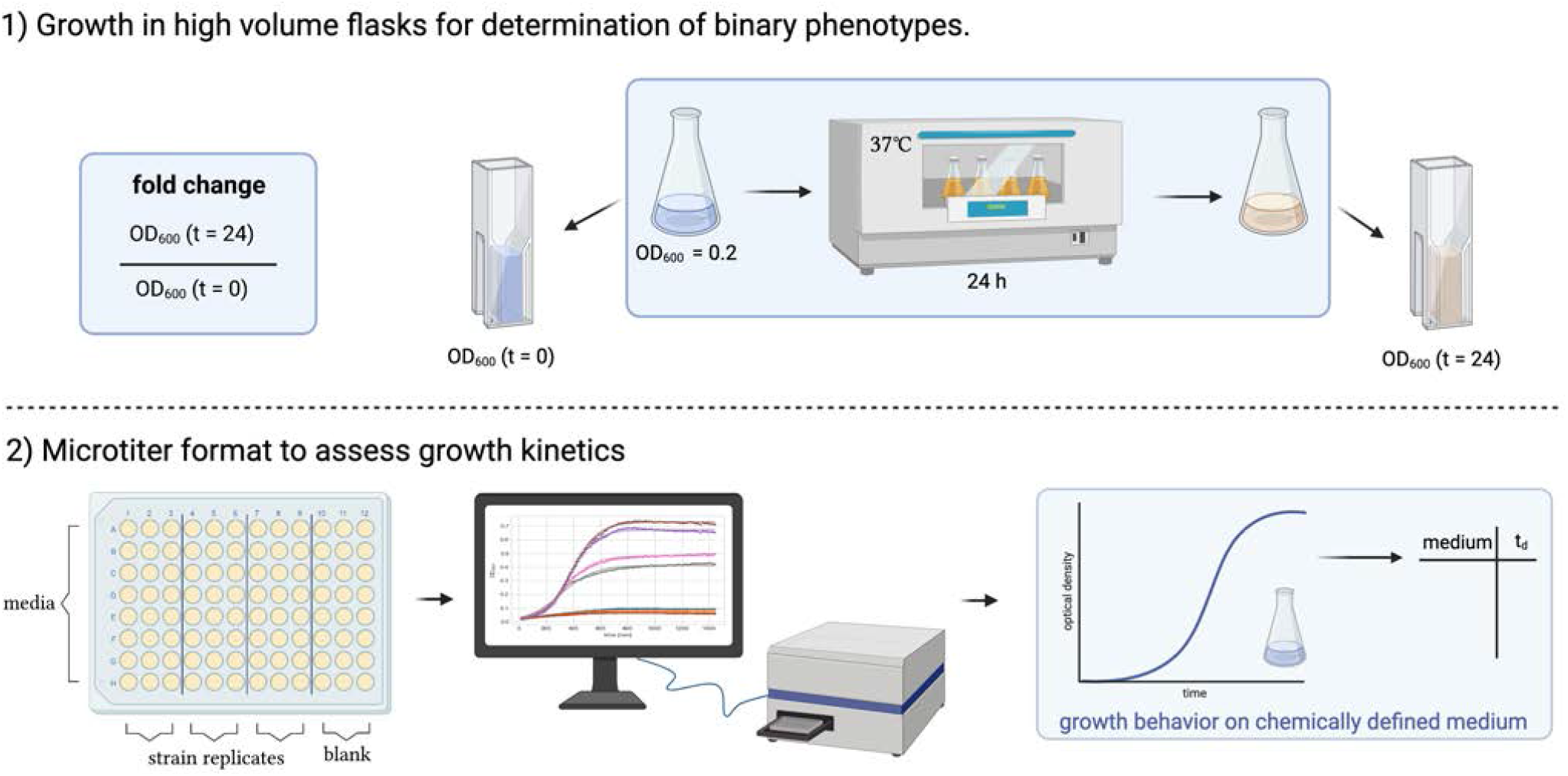
**Graphical summary of two experimental approaches** to study the growth of *C. striatum* under defined nutritional conditions. 1) Growth in high Volume Flasks for binary assessment of growth. 2) Growth in microtiter plate format to assess growth kinetics. The figure was created using BioRender.com.

1. **Growth in higher volume flasks for determination of binary phenotypes**. Higher volumes of media (10 mL in 50 mL flasks) were inoculated to an optical density of 0.2, and the optical density was assessed after 0 h and 24 h to determine binary growth phenotypes (growth vs. no growth). Binary phenotypes are used to assess the accuracy of a GEM where a simulation can be run, e.g., on different carbon sources that can then be tested in the laboratory.
2. **Microtiter format to assess growth kinetics**. Besides binary phenotypes, the prediction of metabolites or additional nutrients that increase/decrease growth rates is of high relevance for the planning of *in vitro* experiments. We used a volume of 150 µL per medium in a 96-well plate and automated the assessment of OD_600_ over time to determine growth kinetics. From the growth curves captured via a plate reader the doubling time at the inflection point of the sigmoidal curve can be extracted and compared to those predicted by the GEM.

#### 3.2.2 *Growth in LB medium is predicted correctly, and growth rates for three strains in LB medium* in silico *are reflected* in vitro

GEM analysis predicted binary growth of TS, 1197, and 1115 while growth of strain 1116 was not possible under the modeled nutritional composition of LB medium (Table 2). Investigation of this phenomenon revealed that strain 1116 lacks a part of the nicotinate metabolism (Figure S2) making the strain auxotrophic for the essential enzymatic cofactor nicotinamide `D`-ribonucleotide (nmn). A putative nmn transporter is predicted by the GEM, supporting the existence of an auxotrophy to this compound. In contrast, transporters for the precursors of nmn (nicotinate or nicotinamide) were not predicted. This caused the GEM to predict a specific need for nmn which was not available in the *in silico* formulation of LB. Interestingly, all strains were able to grow in LB in *in vitro* experiments (Figure 3A). This shows that the auxotrophy of 1116 is not relevant in LB. This might have two reasons. Either nmn might be present in sufficient amounts in LB medium, or its precursors (nicotinate, nicotinamide) are present and taken up by unidentified transport systems in *C. striatum*. Growth rate analysis using microtiter plates and automated recording of the optical densities showed differences between the strains. In LB medium, strain 1197 grew significantly slower (81.13 min*/*Generation(Gen)) than strains 1115 and 1116 (45.92 min*/*Gen and 40.38 min*/*Gen respectively) (Figure 3B). These differences were not predicted by the respective GEMs, suggesting differences in the metabolisms of the strains that are currently not reflected within the models. Strain 1116 differed from strain 1115 in the reached ODs of around 0.2 and 0.6, respectively, but not in the doubling times of 40 to 45 minutes. Only for strain 1197, the doubling times extracted from the growth curves differed notably from those predicted by the model. This can be explained by the experimental data used to determine the *in vitro* doubling time. Strains 1197 and 1116 only grew to an OD of around 0.2 in LB medium, hindering the accurate and reliable fitting of the logistic equation to the growth curve (Figure 3B).

**Table 2.**
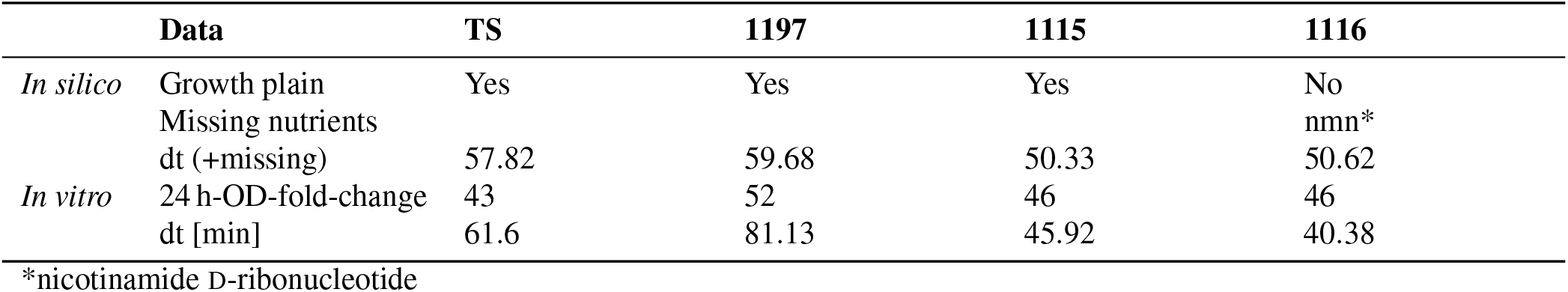
Growth behavior *in silico* and *in vitro* of all strains in LB.

**Figure 3.**
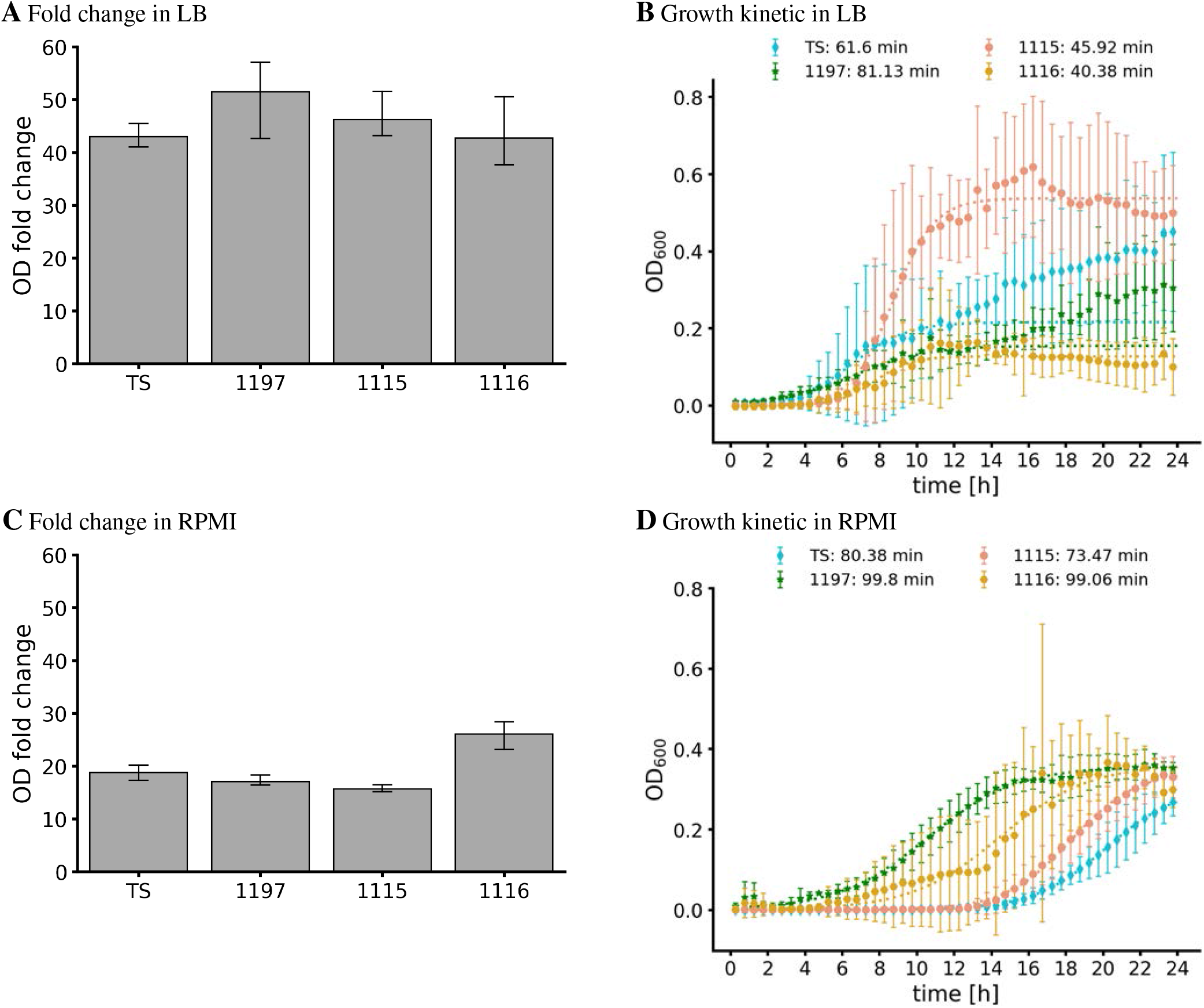
**OD fold changes and *in vitro* growth curves in LB (A, B) and RPMI (C, D)** of the strains TS, 1197, 1115, and 1116. A and C: 10 mL medium in 50 mL flasks were inoculated to OD_600_ = 0.2. After 24 h of incubation, the OD_600_ was measured again, and the fold-change was calculated as a measure for bacterial growth. Shown are the mean and SD of three independent experiments. B and D: Growth in Microtiter plates. 150 mL of medium in a 96 well plate were inoculated and OD_600_ was assessed automatically for 24 h using an Epoch2 plate reader. The logistic fit was calculated using `curve_fit` (dotted lines). The doubling times extracted from the logistic fit are indicated in the legend. For strain TS and 1197, we only fitted data up to 12.5 h to avoid fitting to diphasic growth.

#### 3.2.3 *Model optimization allows congruent* in silico *and* in vitro *growth in RPMI medium*

None of the strains showed *in silico* proliferation on the nutritional composition of RPMI (as detailed by the supplier). Investigating this phenomenon showed a lack of the trace elements (Co^2+^, Cu^2+^, Fe^2+^, Mn^2+^ and Zn^2+^) for all strains while 1116 lacked additionally nmn as observed before. However, all strains grew, effectively in flasks *in vitro* (Figure 3C). This shows the presence of sufficient amounts of trace elements in the medium to allow bacterial growth. Consequently, modification of the *in silico* composition of RPMI by the addition of trace elements is needed to optimize the congruency of *in silico* and *in vitro* analysis. Furthermore, RPMI contains nicotinamide. This strongly suggests that nicotinamide can be taken up by strain 1116 and enables synthesis of nmn.

Growth curve analysis resulted in final optical densities of 0.4 for all strains. However, the curves were characterized by long lag phases and did not show sigmoidal appearances, complicating the extraction of the doubling times. Nevertheless, doubling times ranged from 73 min (strain 1115) to 100 min (strain 1197; Figure 3D). All strains grew slower than predicted (Table 3).

**Table 3.**
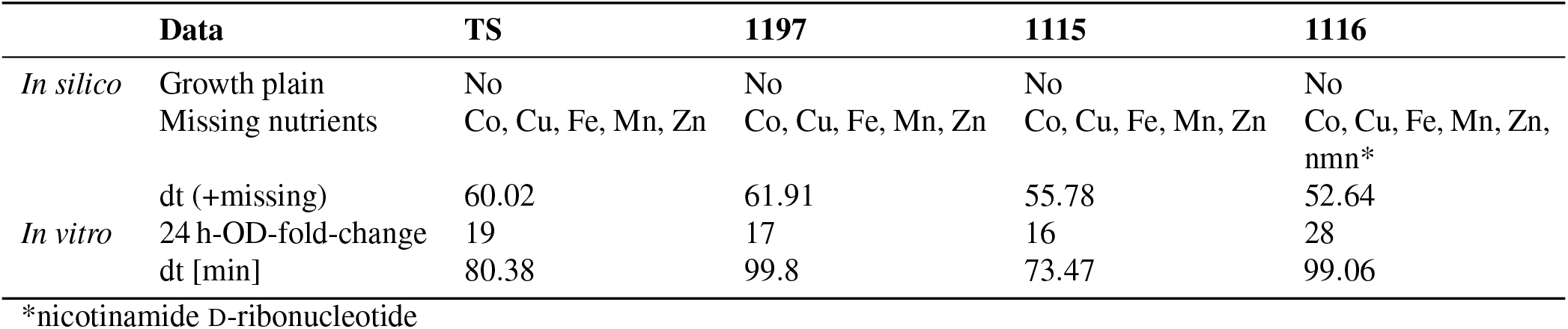
Growth behavior in silico and in vitro of all strains in RPMI. Chemical abbreviations are used for metals.

#### 3.2.4 Growth in CGXII medium requires enrichment to allow

CGXII medium is used as an optimal, chemically defined medium to grow *Corynebacterium glutamicum*^55^. Therefore, we decided to test the ability of our strains to proliferate in this medium. *In silico*, none of the strains were predicted to grow in standard CGXII composition. Analysis of this phenomenon revealed that all strains need the addition of the trace metal cobalt. This allowed *in silico* the proliferation of the strain TS while all other strains showed auxotrophies for `L`-Cysteine and pantothenic acid (strain 1197 and 1115) or nmn and pantothenic acid (strain 1116). The addition of the respective nutrients *in silico* allowed simulated proliferation for all strains (see Table 4). Interestingly, none of the strains, not even the strain TS showed proliferation in CGXII (supplemented with cobalt) containing flasks *in vitro*. As the reasons for this discrepancy were unclear, we tested if the addition of other nutrients might stimulate growth. Firstly, we added 0.2 % Tween80 (Tw) which is known to have growth stimulatory effects on *Corynebacteria*^57^. This increased the reached OD after 24 h slightly (Figure 4A). However, we observed strongly improved growth of strain TS when Tween80 as well as a complex mixture of amino acids (casamino acids – CasA) was added to CGXII.

**Table 4.**
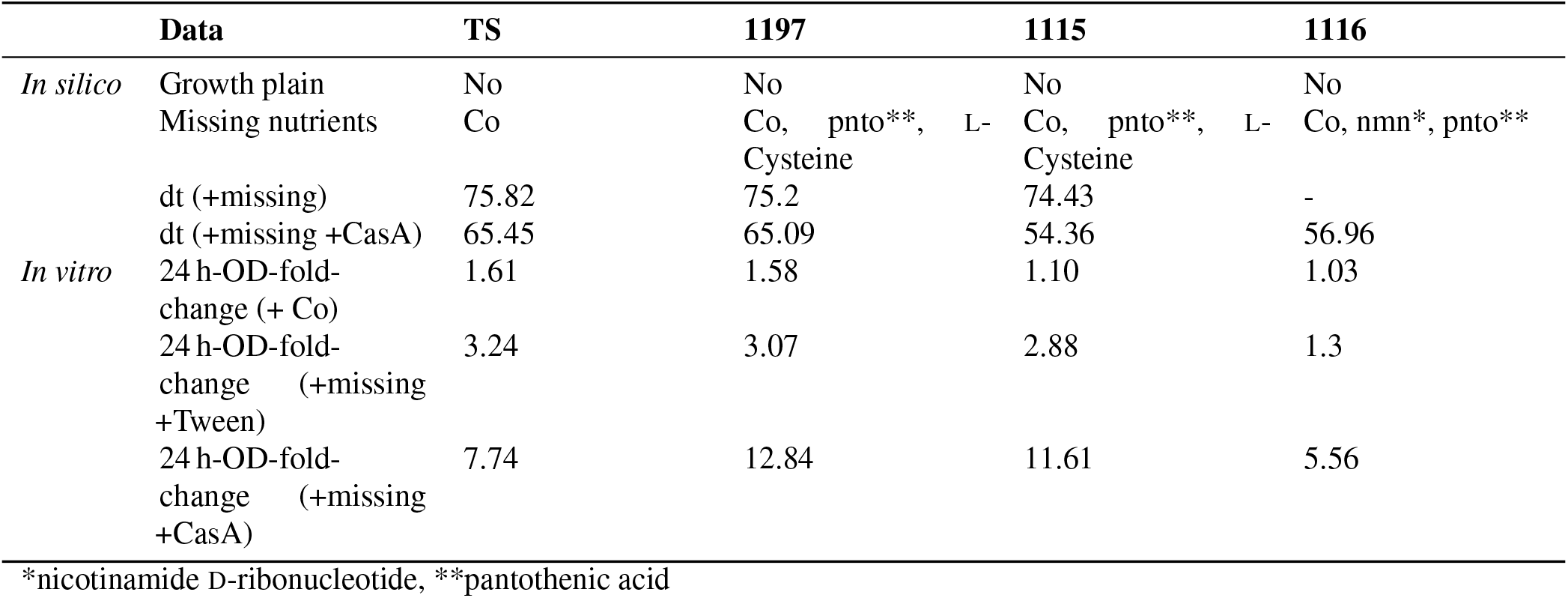
Growth behavior *in silico* and *in vitro* of all strains in CGXII. “-” indicates doubling time is equal to infinity due to zero growth rate. Symbols are used for metals.

**Figure 4.**
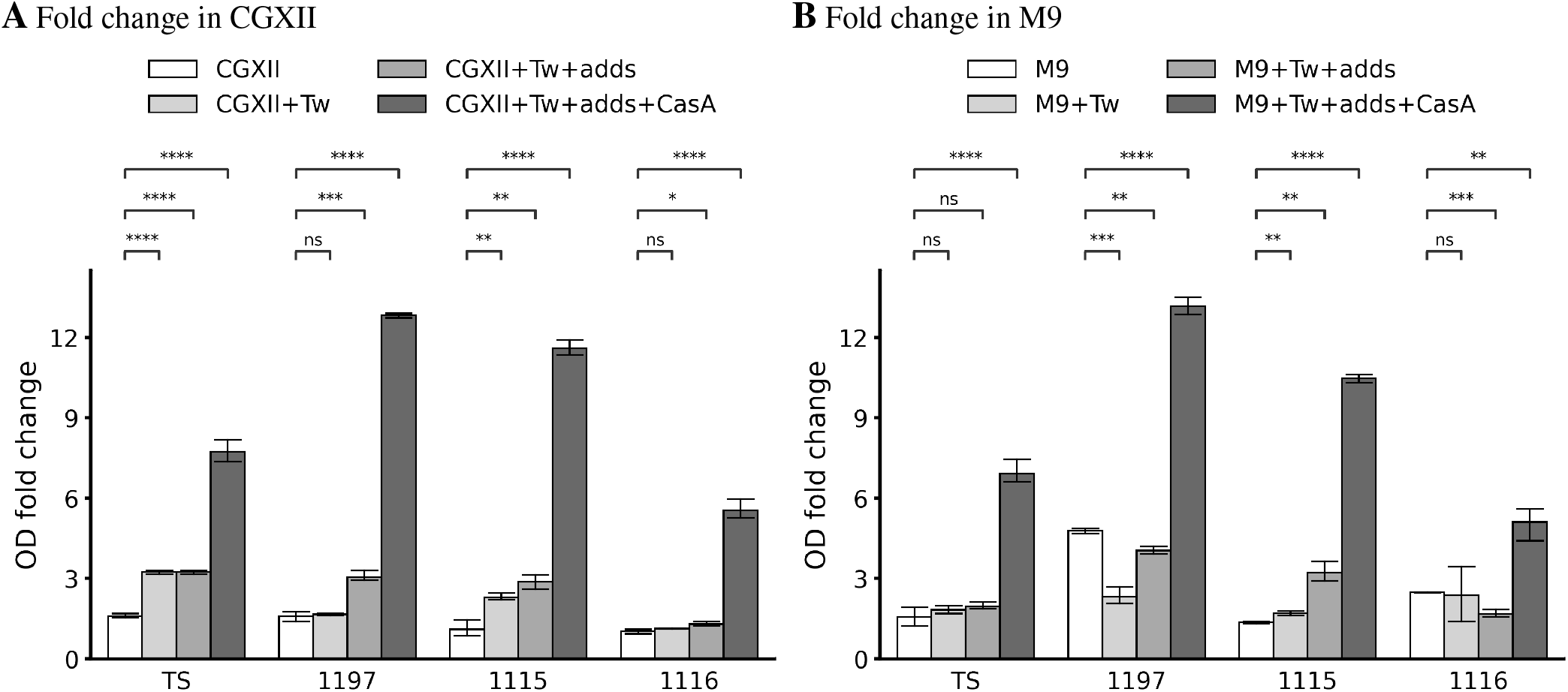
***In vitro* binary growth phenotypes in CGXII (A) and M9 (B)** of the strains TS, 1197, 1115, and 1116. 10 mL medium in 50 mL flasks were inoculated to OD_600_ = 0.2. After 24 h of incubation, the OD_600_ was measured again, and the fold-change was calculated as a measure for bacterial growth. Shown are the mean and SD of three independent experiments. “Tw” corresponds to the addition of 0.2 % Tween 80, “adds” indicates the addition of predicted missing metabolites (see Table 5 and Table 4), “CasA” indicates that 0.1 % casamino acids were added. The concentrations of the added predicted missing metabolites are denoted in Table S1.

Similarly, all other strains failed to grow in cobalt supplemented CGXII and the addition of Tween 80 in combination with strain-specific metabolites (see Table 4) allowed only minor growth improvement. Interestingly, the addition of CasA improved the growth of all strains, but the growth of strain 1116 was improved to a lesser extent compared to the other strains.

None of the strains showed growth in microtiter plate format in CGXII, a phenotype that did not change upon the addition of further nutrients as described above. Accordingly, the growth kinetics of the different strains in CGXII could not be assessed.

#### 3.2.5 Growth in M9 medium requires enrichment to allow proliferation in vitro

M9 is a widely used minimal medium. For growth in M9 our models predicted that all strains needed the addition of trace metals as well as strain-dependent metabolites such as `L`-Cysteine, nicotinamide `D`-ribonucleotide and pantothenic acid (Table 5).

**Table 5.**
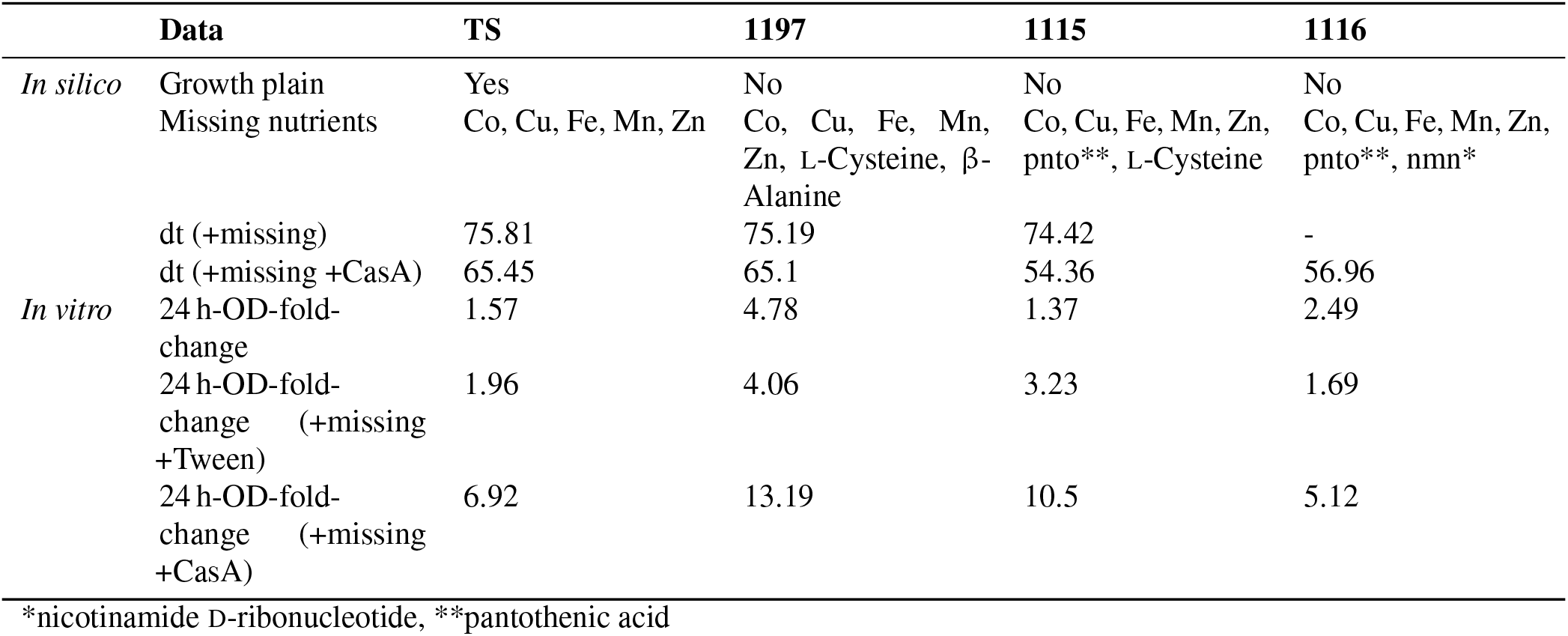
Growth behavior *in silico* and *in vitro* of all strains in M9. “-” indicates doubling time is equal to infinity due to zero growth rate. For *in vitro* experiments β-Alanine was replaced with Pantothenic acid. Symbols are used for metals.

Again, the *in vitro* experiments did not reflect the *in silico* prediction. Growth in M9 medium (supplemented with all trace metals) was poor for all strains. The addition of Tw as well as of the respective “missing” metabolites did only improve the growth for all strains but the addition of CasA increased the proliferation. Similar to growth in CGXII, we failed to detect growth in M9 in microtiter plate format which prevented growth rate analysis.

### 3.3 RefineGEMs: A toolbox for faster curation and analysis

The work on reconstructing multiple strain-specific models of opportunistic bacterial pathogens resulted in a new, more general software toolbox called refineGEMs. It combines, integrates, and extends COBRApy and the libSBML^58^ Python package for faster and more accessible GEM curation and analysis in a standardized repository structure.

The toolbox offers various features to help in the investigation of any GEM. It enables the user to load GEMs from SBML^19^ files, build a report containing key entities such as charge unbalanced reactions and numbers of reactions, and compare genes present in the model to those found in the KEGG Database given a GFF file and the KEGG ID of the organism. The charges and masses of the metabolites in the model can also be compared to those found in the ModelSEED database^59^. The toolbox facilitates performing reproducible growth simulations.

In addition to investigating GEMs, refineGEMs can also be used to curate a given model. For instance, if a model was created with CarveMe version 1.5.1, refineGEMs can transfer relevant information from the notes field within model components to their respective annotation section and can automatically annotate GeneProducts in the model from their IDs using the NCBI IDs. It also enables the addition of KEGG Pathways as groups (using the SBML^19^ groups extension) and SBO term annotation refinement via its integrated SBOannotator^39^. Other important functionalities are updating the annotation of metabolites and extending the model with missing reactions based on a table filled in by the user during manual research.

RefineGEMs can be used in two different ways: (a) as a standalone script using the 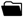 `main.py` script and the corresponding 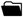 `config.yml` file, which is available on GitHub (b) it can be installed via `pip` and the functions can be accessed individually Comprehensive documentation is available online, which is accessible via GitHub. The scope of the different modules of refineGEMs is outlined in Table 6, giving an overview of the capabilities of the toolbox. All growth simulations and parts of the model refinement described in the results above were done using refineGEMs.

**Table 6.** Overview of refineGEMs modules and their scope. More in-depth information can be found in the documentation accessible via GitHub. The `sboann` module is based on the SBOannotator’s original implementation from the bachelor thesis by Elisabeth Fritze.

| Module | Aim |
| --- | --- |
| analysis_biocyc | All functions required solely for the comparison to BioCyc<sup>60</sup> |
| analysis_db | All functions required for the KEGG and BioCyc comparison |
| analysis_kegg | All functions required solely for the comparison to KEGG<sup>43</sup> |
| charges | Add charges from ModelSEED<sup>59</sup> to non-charged metabolites |
| comparison | Comparison of growth behavior of multiple strains |
| curate | Model refinement, synchronization of annotations, transfer of notes to annotations |
| cvterms | Utilities for CVTerms (f. ex. extraction of different database IDs) |
| databases | Holds functions to access databases used in curation |
| gapfill | Model extension by comparison to BioCyc or BioCyc and KEGG |
| growth | Growth simulation on SNM<sup>61</sup>, LB, M9, SMM, CGXII and RPMI |
| investigate | Model scope and quality assessment |
| io | Utilities to load and write models in XML format |
| modelseed | Comparison of charges and formulae to the ModelSEED database |
| pathways | Addition of KEGG pathways as groups based on the reaction annotations |
| polish | Correction of draft models obtained with CarveMe |
| sboann | Refinement of SBO terms |

## 4 Discussion

This study aimed at building strain-specific genome-scale metabolic models (GEMs) for *Corynebacterium striatum*. Only strains whose genome has been entirely sequenced and available in the German Collection of Microorganisms and Cell Cultures (DSMZ) were selected as strains to be studied. In doing so, this work comprises three principal objectives: First, developing a unified infrastructure for creating systems biology models was and making it publicly available. Second, using this infrastructure to create GEMs for selected strains of the bacterium *C. striatum* and to make them freely accessible. Finally, experimentally verifying and using these strain-specific GEMs for their further refinement based on generated predictions of bacterial growth in the presence of defined nutritional environments.

The GEMs’ prediction of bacterial growth have enormous potential for laboratory-based microbiology as it might help to quickly adjust culture conditions to optimize growth rates and growth yields, to optimize fermentation processes or even to enable growth of so far not-culturable bacteria. However, while GEMs are nowadays created for many different species and even for multiple strains of the same species, their ability to correctly predict biological phenotypes is hardly investigated.

In a typical scenario, published computer models are initially developed as a theoretical foundation and only checked against previously published data, if available^62^. In other cases, previously published models get revised based on recent developments^14, 63^. Other research groups can perform a comprehensive analysis and validation once these models are publicly available. In the case of the coronavirus pandemic, host-virus models initially developed purely theoretically^64^ could be used as the basis for later experimental studies that enabled the identification of potential antiviral agents^65^. However, it is not always possible for researchers working on systems biology modeling to have their models tested in the laboratory, but ideally, this should be standard practice. To close this gap we used a tightly integrated collaboration between bioinformatics and systems biology, on the one hand, and microbiology, on the other hand. We created novel GEMs for several *C. striatum* strains and used them to make predictions about proliferation of the strains under defined nutritional conditions. These predictions were then compared to biological data sets gained in the laboratory. All models are available for download from the BioModels database.

Our experiments identified several pitfalls reducing accuracy of model prediction. We found several examples of “false negative” predictions, referring to *in vitro* growth while GEMs predicted a growth failure. This phenomenon can have two different underlying reasons. Firstly, it is possible that certain nutrients are present *in vitro* that were not included in *in silico* formulation of the nutritional composition. This highlights that the *in silico* assembly of the nutritional composition of laboratory media needs to be approached deliberately. Trace metals are important to mention in this regard. These nutrients are hardly mentioned on ingredient lists of commercial media (e.g., RPMI). Similarly, not all trace metals are purposely added to defined media (CGXII, M9) produced in laboratories around the world. However, the amount needed is extremely low and physiologically relevant concentrations are available in most media as long as special action for their removal (e.g., addition of chelators) is avoided. Accordingly, media formulations used with GEMs should reflect that trace metals are most likely not limiting factors in experimental setups.

Furthermore, for complex media ingredients such as yeast, tryptic soy or meat extracts that are frequently used in microbiological practice, the precise composition of these extracts are unknown. However, they contain excess of amino acids, vitamins, ribonucleotides etc. which should be included in the *in silico* composition of media to increase the accuracy of growth prediction.

Secondly, false negative predictions can also be caused by inappropriate transport activities within GEMs. One example in this regard is nicotinic acid transport. The GEM of strain 1116 predicts nmn to be a crucial metabolite for this strain. Supplementation of the medium with the precursors nicotinic acid or nicotinamide is regarded as insufficient as specific transporters for their import are not predicted within the genome. In contrast to this prediction, we found the strain to proliferate in the presence of nicotinamide (growth in RPMI) as well as in the presence of nicotinic acid (M9 medium), strongly suggesting that both intermediates can be acquired by the bacterium and enable the synthesis of nmn. However, additional experimental evidence is needed to validate this hypothesis and the responsible transport systems remain to be identified.

We also observed “false positive” predictions, referring to GEMs predicting growth, while proliferation is not observed *in vitro*. These inconsistencies are more problematic, as they reduce the usefulness of GEMs for laboratory-based experiments. Unfortunately, the reasons for this are less clear and the issues are more difficult to address. However, one reason might be that GEMs are based on genetic information while information about expression of the respective metabolic pathways is missing. Although genes for biosynthesis of aminoacids or vitamins might be present within a strain of interest, a failure to express the same might entail physiological auxotrophies. This phenomenon is known for the amino acids Leucin and Valin for *Staphylococcus aureus*^66, 67^. Along this line we found that the addition of casamino acids improved proliferation of *C. striatum* in CGXII and M9 media.

Additionally, *in silico* assumptions are frequently not directly transferable to *in vitro* conditions. GEMs assume homogeneously mixed cells under isothermal and isobaric conditions whose compartments maintain constant volumes. Molecular concentrations are assumed abundant enough to be effectively continuous. In addition, for simplicity, some physicochemical factors, such as osmotic pressure or electroneutrality, are not considered. The modeled cells would also remain steady, resulting in constant molecular concentrations. These assumptions are by far not met *in vitro* which can also account for discrepancies between model experimental results. Taken together, GEMs have high potential to support experimental microbiologists. However, close interactions and repetitive cycles of prediction experimental validation and model refinements are needed to improve accuracy of the predictions.

Inclusion of experimental observations into GEMs represents another task for the future. Exemplarily, it is known that addition of Tween80 improves *in vitro* growth of *Corynebacteria*. It is currently impossible to incorporate such findings into GEMs.

Since all individual steps for reconstructing these five strain-specific *C. striatum* models were performed by standardized Python scripts in a directory structure based on version control and made available, the modeling process can be traced in detail. Thus, a software infrastructure comprehensively tested on several genuine and relevant case studies is available, which can be applied to further reconstruction projects.

## 5 Conclusions

The tools and models created in this work through close collaboration between the dry and wet laboratories provide a valuable working basis for subsequent studies. The software infrastructure developed and the models created have been extensively tested and validated using standard methods.

However, since no computer model, no matter how carefully developed, can ever exhaustively represent all processes of a cell type under investigation, subsequent work will be necessary to elucidate further the metabolism of all five *C. striatum* strains described herein in order to provide even more accurate and thus more meaningful predictions.

Concerning the underlying model assumptions, extensions for gene expression would also be beneficial. Modeling approaches are already available as ME models (for Metabolism and Expression^68^). On the other hand, it may be interesting to shed light on the effects of single gene mutations on enzyme efficiency and, thus, metabolism as a whole, for which so-called GEM-PRO models could be used^69^. This would be particularly important for clinical isolates. With the increasing availability of genome sequencing, individual patient germs could be characterized.

Further developments on the experimental side would also be beneficial: since the predictive power improves the more precisely the media composition is known, the substance concentrations of their components should be clarified more precisely for model development using human microbiota. Due to biovariability, only average values can be expected, but knowledge of the standard deviation across subjects also allows conclusions to be drawn and enables computer experiments under variation with higher accuracy.

Since no biological system can live in isolation, the interaction with other members of the microbiota remains an essential aspect for subsequent research to understand the diverse interactions with commensals, other pathogens, and the human host in a larger picture. Basic approaches in this direction are already available, with which the models developed here can also be combined^70–72^.

The models, Python programs, and Git-based version management working template presented with this work can be directly applied and used for subsequent projects immediately and can be independently further developed. Thus, an infrastructure extensively tested on several relevant reconstruction efforts is available and can be applied to any GEM.

## Author contributions

F. B. created and curated the models, developed the binary growth phenotype experimental protocol, conducted the experiments and introduced the `refineGEMs` toolbox. G. O. G. extended the `refineGEMs` toolbox. L. C. supervised the laboratory work. S. H. supervised the *in vitro* study. A. D. supervised the *in silico* study. F. B., S. H., and A. D. wrote the manuscript, which all authors have read and approved.

## Acknowledgments

The authors thank Libera Lo Presti for manuscript reviewing and proof-reading.

## Data availability statement

All models are available on BioModels and were submitted according to the standards set in the community to enable the curation of those models within the database. The models were submitted in a Computational Modeling in Biology Network (COMBINE) archive Open Modeling EXchange format (OMEX) format including the respective MEMOTE and Flux Variability Analysis (FVA), Reaction deletion, Objective function values, Gene deletion fluxes (FROG) reports. Annotations of all models were formatted adhering to community standards. All models are also available via GitHub where future changes and additions will be continuously implemented. The GitHub repository was also used during GEM reconstruction and thus holds previous versions of the models. Its structure is based on the standard-GEM format. The MEMOTE diff report can be found on the GitHub repository as well.

## Supporting Information

### Media preparation

LB (Sigma), BHI (Roth), TSB (Oxoid), and RPMI (Thermo Fischer^51^) powders were dissolved in MilliQ water. Afterward, the media were sterilized by autoclaving or by 0.22 µm filtration for RPMI.

M9 Minimal Salts were ordered from Sigma (M6030)^53^. The M9 medium was then prepared using a 5× autoclaved stock of the M9 salts with the addition of 1 mM MgSO_4_ · 7 H_2_O, 0.1 mM CaCl_2_ · 2 H_2_O, 0.1 % `D`-glucose which were all sterilized by filtering through a 0.22 µm filter.

CGXII was prepared according to the protocol given by Unthan et al. 55. All components were prepared as stock solutions in water, and sterilized by filtering through a 0.22 µm filter. The final medium was then prepared by mixing the stock solutions, adding water to fill to the desired volume, and adjusting the pH to 7 using a 1 M NaOH solution. Afterward, the medium was sterilized by filtering.

All tested additives were prepared as stock solutions, sterilized by filtering, and then added to obtain the desired concentration as shown in Table S1. The trace element solution contains MnCl_2_ · 4 H_2_O, ZnCl_2_, CuCl_2_ · 2 H_2_O, CoCl_2_ · 6 H_2_O, Na_2_MoO_4_ · 2 H_2_O, H_3_BO_3_ and NiCl_2_ · 6 H_2_O.

**Table S1.** Metabolites that were used as additives for growth enhancement. This table holds the concentration that was used if the metabolite was added and the paper that this concentration was taken from.

| metabolite | concentration [g/L] | based on |
| --- | --- | --- |
| Nicotinate | 0.001 | Yao et al. (2018) |
| (R)-Pantothenate | 0.001 | Yao et al. (2018) |
| Co<sup>2+</sup> | 0.0024 | Taylor and Holland (1989) |
| L-Cysteine | 0.05 | Concentration in RPMI<sup>51</sup> |
| Fe<sup>2+</sup> | 0.01 | Concentration in CGXII<sup>55</sup> |

### Growth in liquid medium

All doubling times were extracted by fitting a logistic equation to the growth data. Growth curves with extracted doubling times in TSB are shown in Figure S1.

**Figure S1.**
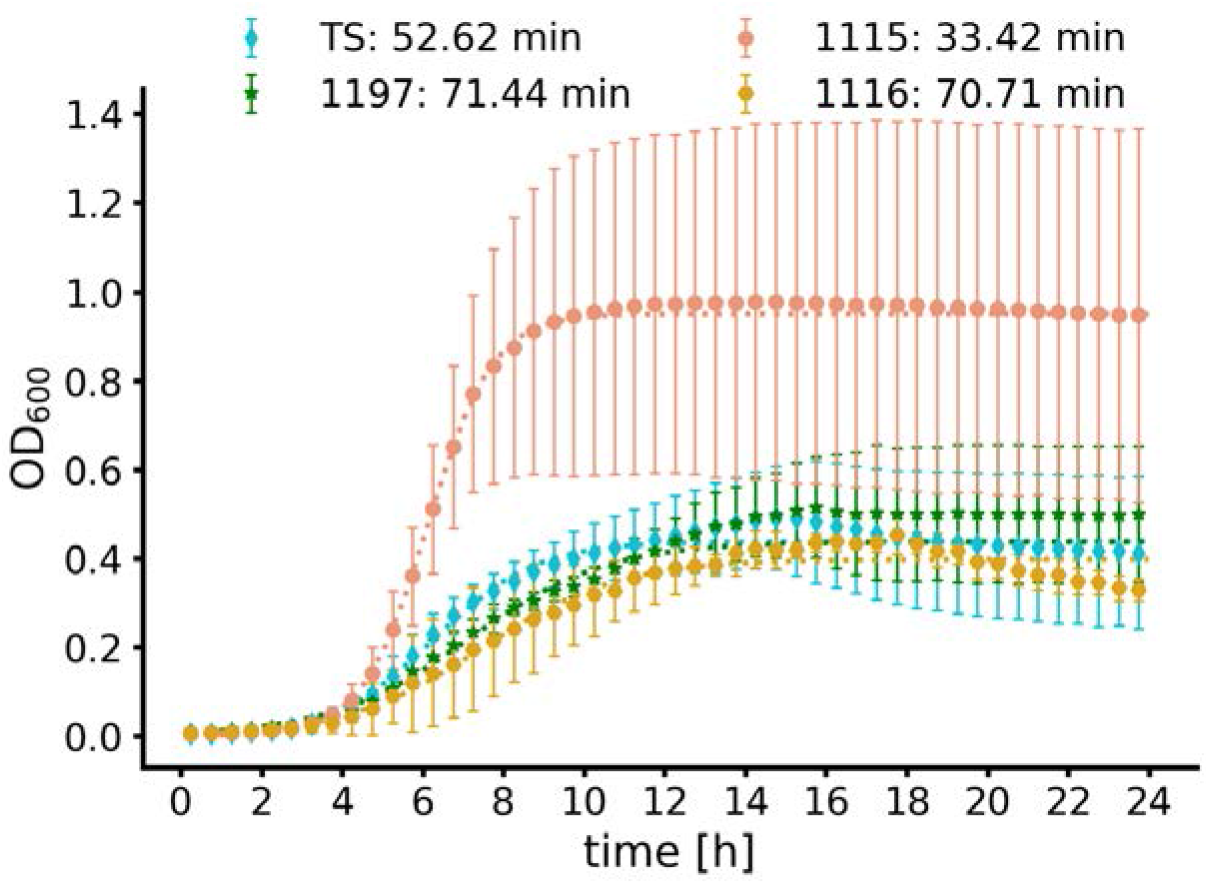
*In vitro* growth curve in TSB of the strains TS, 1197, 1115 and 1116. Growth *in vitro* in TSB was determined via OD600 measurements for 24 h (data points). Shown are the means of *n* = 3 independent experiments with standard deviation. Logistic fit was then calculated using `curve_fit` (dotted lines). The doubling times extracted from the logistic fit are indicated in the legend. For strain TS and 1197 we only fitted data up to 12.5h to avoid fitting to diphasic growth.

### Model status

**Table S2.** Overview on key numbers of all models. Strains TS, 1197, 1115 and 1116 were compared to laboratory work. Strain KC was not available for laboratory experiments.

| strain | TS | 1197 | 1115 | 1116 | KC |
| --- | --- | --- | --- | --- | --- |
| # metabolites | 1,382 | 1,141 | 1,197 | 1,212 | 1,053 |
| # reactions | 2,002 | 1,674 | 1,750 | 1,788 | 1,541 |
| # genes | 772 | 756 | 763 | 755 | 719 |
| metabolic coverage [in %] | 2.59 | 2.21 | 2.29 | 2.37 | 2.14 |
| MEMOTE score [in %] | 86.4 | 86.7 | 86.2 | 86.3 | 84.1 |

### *In silico* media composition

**Table S3.**
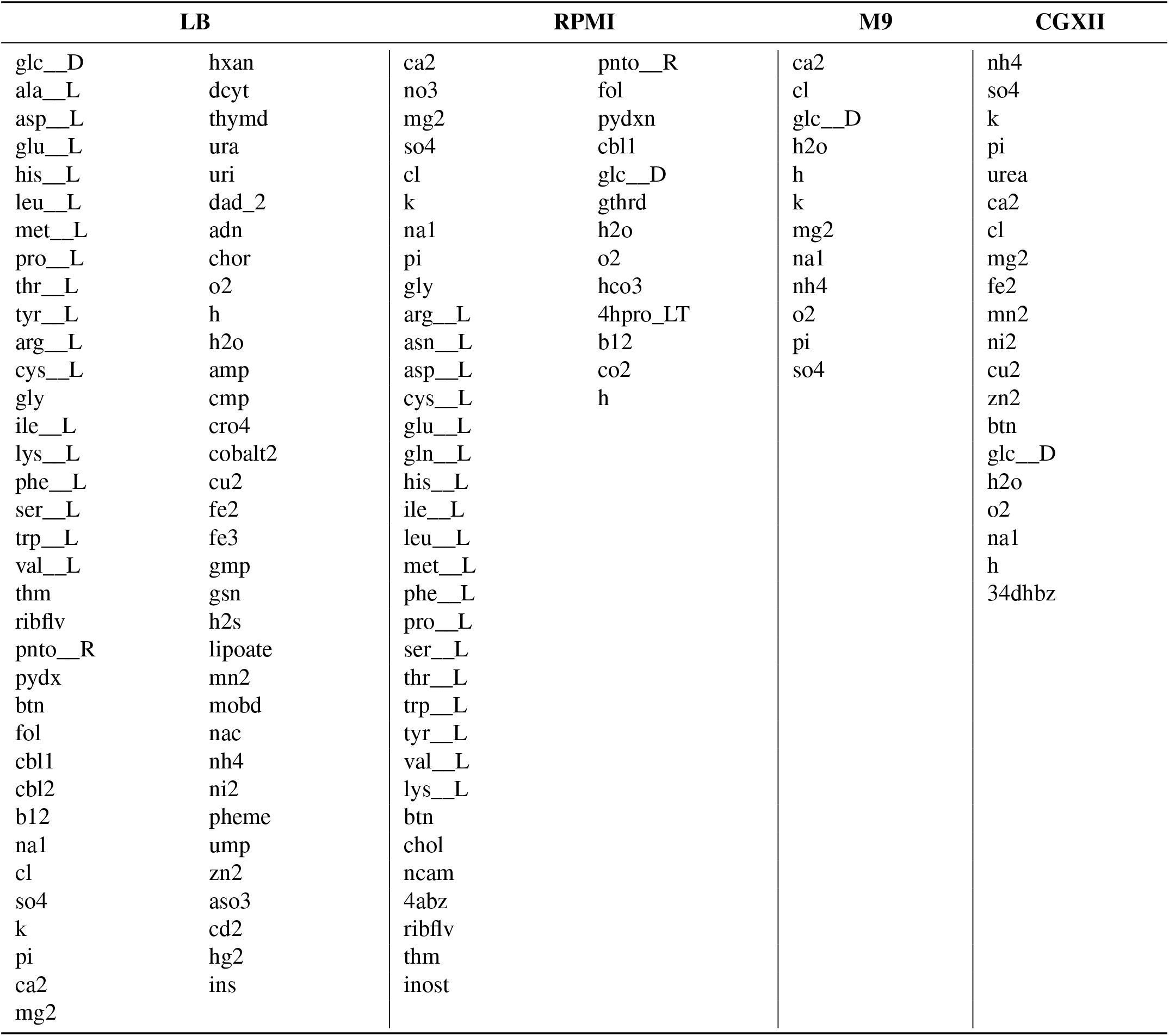
*In silico* composition of the different media used in this study. Indicated are the BiGG IDs of the metabolites for which exchanges were opened during simulation.

### Escher map of strain 1116

**Figure S2.**
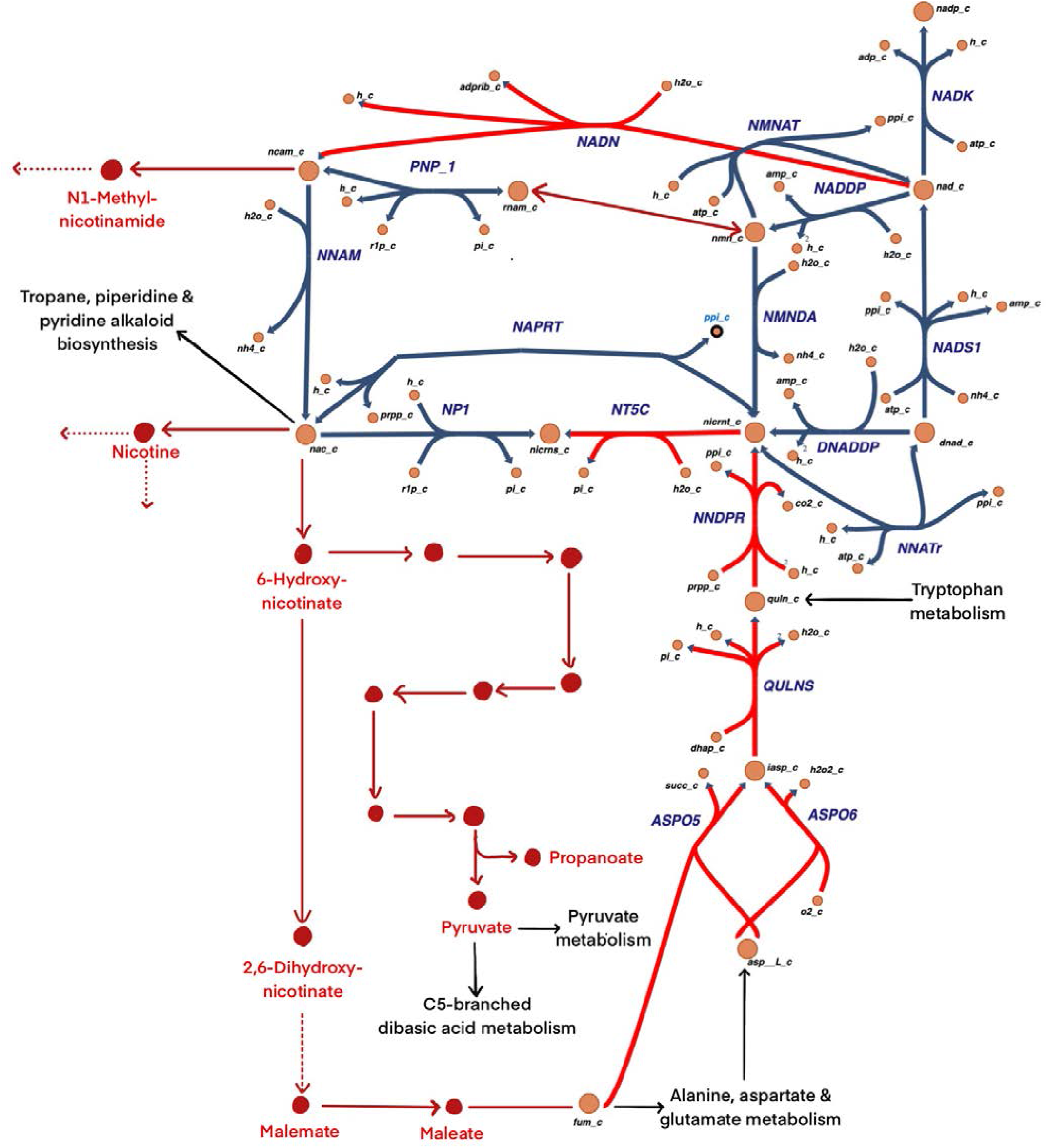
Escher^24^ map of the nicotinate metabolism. Links to other pathways are shown on the map in black. Missing reactions from strain 1116 are shown in red.

